# Epithelial IL-2 is critical for NK cell-mediated cancer immunosurveillance in mammary glands

**DOI:** 10.1101/2024.04.25.591178

**Authors:** Lei Wang, Chandra K Maharjan, Nicholas Borcherding, Rohan P Master, Jiao Mo, Tanzia Islam Tithi, Myung-Chul Kim, Madison E Carelock, Anuj P. Master, Katherine N. Gibson-Corley, Ryan H. Kolb, Kendall A. Smith, Weizhou Zhang

## Abstract

Interleukin 2 (IL-2) is the first identified cytokine and its interaction with receptors has been known to shape the immune responses in many lymphoid or non-lymphoid tissues for more than four decades. Active T cells are the primary cellular source for IL-2 production and epithelial cells have never been considered the major cellular source of IL-2 under physiological conditions. It is, however, tempting to speculate that epithelial cells could potentially express IL-2 that regulates the intricate interactions between epithelial cells and lymphocytes. Datamining our recently published single-cell RNAseq in the mouse mammary gland identified IL-2 expression in mammary epithelial cells, which is induced by prolactin via the STAT5 signaling pathway. Furthermore, epithelial IL-2 plays a crucial role in maintaining the physiological functions of natural killer (NK) cells within the mammary glands. IL-2 deletion in the mammary epithelial cells leads to a significant reduction in the number and function of NK cells, which in turn results in defective immunosurveillance, expansion of luminal epithelial cells, and tumor development. Interestingly, T cells in the mammary glands are not changed, indicating the specific regulation of NK cells by epithelial IL-2 production. In agreement, we also found that human epithelial cells express IL-2 and NK cells express the highest level of IL2RB among all the immune cells. Here, we provide the first evidence that epithelial cells produce IL-2, which is critical for maintaining the physiological functions of NK cells in immunosurveillance.

## Introduction

Interleukin-2 (IL-2) was identified as a T cell growth factor four decades ago and plays multifaceted roles in immune regulation by interacting with IL-2 receptors (IL-2Rs) across various immune cells^1–3^. Regulatory T cells (Tregs) are the major effector cell type of IL-2 and are crucial for ensuring immune homeostasis^4, 5^. Deficiencies in IL-2 or IL-2R in mouse models lead to severe autoimmunity or immunodeficiencies, highlighting the pivotal role of IL-2 in immune tolerance^6–8^. IL-2 signaling differentially influences T cell fate, with a high level of IL-2 signaling essential for effector T cell activation, while a low level for the transition to memory T cells^9^. IL-2 also significantly impacts natural killer (NK) cell proliferation and cytotoxicity, underscoring its impact on innate immunosurveillance^2, 10^. IL-2 is predominantly produced by active T cells and recent discoveries have expanded cellular sources for IL-2 production, such as dendritic cells (DCs) and group 3 innate lymphoid cells (ILC3) in the intestine^11–13^. The different cellular sources of IL-2 presumably provide spatially and temporally controlled regulation of its effector cells^13–17^. To the best of our knowledge, there is no evidence showing that epithelial cells produce IL-2 under physiological conditions and whether epithelial IL-2 plays any physiological role in immune homeostasis within tissues.

The immune landscape within the mammary gland mirrors that of mucosal organs, with both adaptive and innate immune cells orchestrating physiological processes such as immunosurveillance, tolerance, and defense against inflammation and oncogenesis. Innate immune cells, such as eosinophils, mast cells, and macrophages, have been shown to regulate mammary morphogenesis^18–21^. T helper type 1 (T_h_1) cells, however, inhibit ductal morphogenesis of the mammary glands through interferon-gamma (IFN-γ) secretion^22^. There is not much evidence of how mammary epithelial cells shape the local immune homeostasis.

The current study unveils a critical role of epithelial cell IL-2 in NK cell function and immunosurveillance. IL-2 deletion in mammary epithelial cells leads to an increased tumorigenesis rate in aged mice, associated with the reduction of NK cell number and activity. Our findings uncover an unanticipated source of IL-2, shedding light on its integral role in the immunosurveillance against tumorigenesis.

## Results

### Mammary epithelial cells (MEC) express interleukin-2 (IL-2)

We have performed a single-cell RNA sequencing (scRNA-seq) study to analyze MECs from virgin mice^23^, where we identified 15 distinct MEC clusters (Fig. 1A, left panel, reanalyzed from GSE143159) using the uniform manifold approximation and projection (UMAP) method. These clusters were further identified as luminal progenitor cells (LPC), mature luminal cells (LMC), basal cells (BC), and protein C receptor-positive (PROCR^+^) stem cells (PSC) (Fig. 1A, left panel). The expression of *Il2* mRNA across these MEC subtypes was noted in all 15 clusters (Fig. 1A, right panel), with *Il2* positivity ranging from 2-3% to 20% (Fig. 1B). We isolated single cells from the fourth mammary gland (MG) after removing the inguinal lymph node. Following 4-hour cell activation cocktail (w/ Brefeldin A) administration, we assessed IL-2 protein expression in T cells (CD45^+^CD3^+^), stromal cells (CD45^-^CD31^-^Ter119^-^CD24^-^CD49^-^), luminal (CD45^-^ CD24^+^CD49^low^, LMECs), and basal (CD45^-^CD24^low^CD49^+^, BMECs) epithelial cells (sFig. 1A, gating scheme). IL-2 expression was detected in both LMEC and BMEC, to a similar extent as active T cells (Fig. 1C-1D). Stromal cells – most likely fibroblasts in the MG, did not exhibit any expression of IL-2 protein (Fig. 1C-1D). Using a bulk RNAseq dataset including the most comprehensive mouse tissues with at least two replicates per tissue type^24^, we found that mouse MG expresses the highest level of *Il2* mRNA, even higher than that from lymphoid tissues such as the thymus and spleen (Fig. 1E). Furthermore, the HC11 cell line, derived from BALB/c mouse mammary epithelium, was able to differentiate into LMECs under the culture medium containing fetal calf serum, supplemented hydrocortisone, insulin and prolactin (HIP) (sFig. 1B). IL-2 protein was detectable in HC11 cells cultured with either control medium or HIP medium, with HIP medium showing slightly elevated level of IL-2 protein (sFig. 1C-D), in agreement with the LMECs expressing higher level of IL-2. In conclusion, our data identified the expression of IL-2 at mRNA and/or protein levels in both LMECs and BMECs.

**Figure 1.**
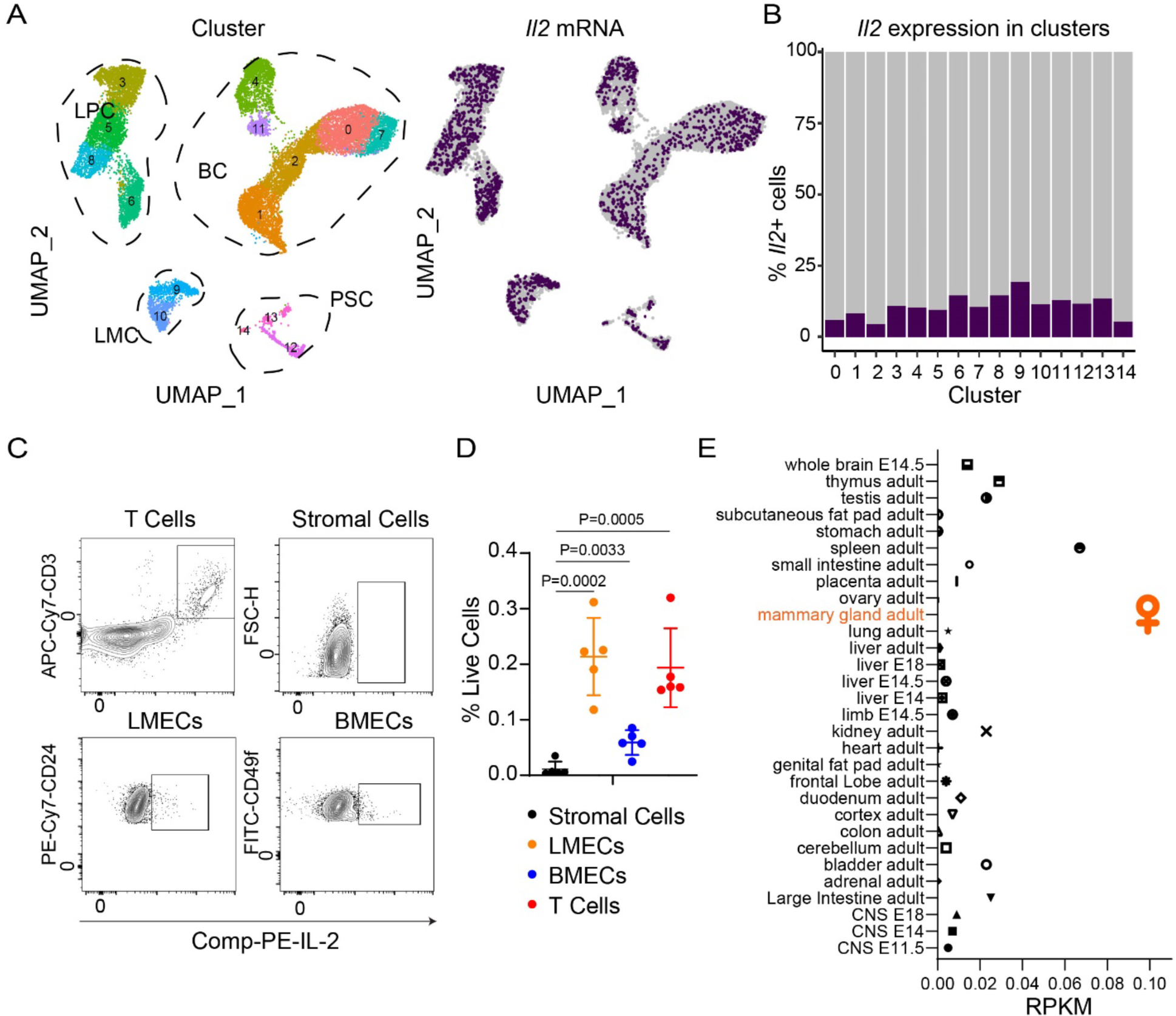
MECs express *Il2* in mouse models. **A.** *Il2* mRNA level in clusters of MECs. UMAP of multiple clusters of MECs from scRNA-seq analysis (left) and *Il2* expression in different clusters (Right). LPC - luminal progenitor cells (clusters 3,5,6,8), LMC - mature luminal cells (clusters 9-10), BC - basal cells (clusters 0,1,2,4,7 and 11), and PSC - protein C receptor-positive (PROCR^+^) stem cells (clusters 12-14). **B.** The proportion of *Il2* expression in 15 clusters. **C.** The schematic of IL-2 positive populations gating for flow cytometry analysis, including T, stromal, luminal and basal cells. **D.** Percentage of *Il2* positive cell population of live cells. A two-sided unpaired *t* test was performed, with P values indicated, NS is non-significant. **E.** *Il2* mRNA expression across all mouse tissues – adapted from Mouse ENCODE transcriptome data (BioProject: PRJNA66167).

*Il2* mRNA was readily detectable in human female mammary glands in bulk RNAseq (sFig. 2A), with single-cell expression data expression echoing the detectable *Il2* mRNA in human MECs (sFig. 2B). Several human breast cancer cell lines exhibited various but detectable *Il2* mRNA as well (sFig. 2C).

### Prolactin induces IL-2 expression in mammary cells

IL-2 expression in active T cells has been well-documented and involves a collective activation of transcriptional factors such as NFAT, AP-1, OCT-1, and NF-κB families^25, 26^. To unravel the mechanisms governing IL-2 expression within the MECs, we used a human breast cancer cell line T47D that is known to express estrogen receptor (ER), progesterone receptor (PR) and prolactin receptors and has detectable *Il2 mRNA* expression (sFig. 2C). Since many signaling pathways within the mammary glands can activate similar transcriptional factors as in active T cells, we treated T47D cells with various stimuli including estrogen (E2), progesterone (P4), prolactin (PRL), WNT5A, and/or RANKL, among which prolactin induced significant *Il2* mRNA (Fig. 2A). In contrast, estrogen and progesterone slightly inhibited its expression (Fig. 2A). Wnt5A and RANKL, two other cytokine/growth factors involved in mammary gland morphogenesis or breast cancer^27–29^, inhibit the expression level of *Il2* (sFig. 2D). In addition, we induced differentiation in mouse immortalized HC11 mammary cells^30^ using HIP medium including prolactin that induced a significant elevation in *Il2* expression (sFig. 1C).

**Figure 2.**
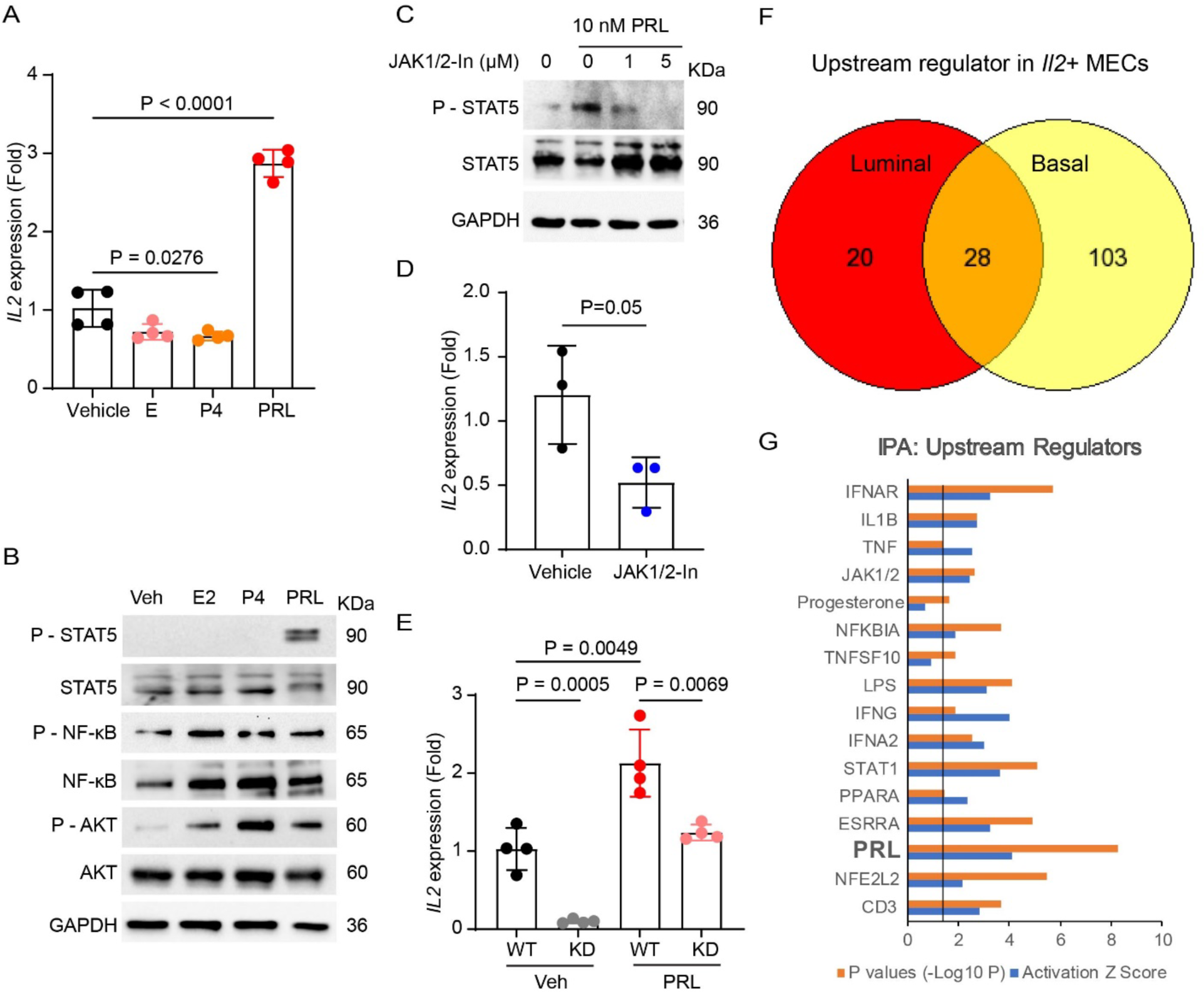
Prolactin induces *Il2* expression. **A.** The fold-change of *Il2* expression in T47D cell line with 48 hours, 10 nM of estrogen (E), progesterone (P4), or prolactin (PRL). **B.** Signaling phosphorylation after 30 mins, 10 nM E, P4 or PRL treatment. Western Blot of phosphorylation of STAT5, NK-κ, AKT. **C.** STAT5 phosphorylation inhibition with JAK 1/2 inhibitor treatment. **D.** *Il-2* expression was inhibited by the JAK 1/2 inhibitor in the T47D cell line. **E.** *Il-2* expression was significantly reduced in STAT5 KD T47D cell line in both vehicle and PRL treatment. **F-G.** Upstream regulators of *Il2+* LMECs and BMECs. **(F)** The Venn diagram showing overlapping upstream regulators comparing *Il2+* and *Il2*-populations. **(G)**. The listed genes of upstream regulators elevated the *Il2+* MECs. A two-sided unpaired *t* test was performed, with P values indicated, NS is non-significant.

When T47D cells were treated with PRL, a plethora of signaling events were induced, including the phosphorylation of Signal Transducer and Activator of Transcription 5 (STAT5), AKT, as well as the induction of NF-κB (Fig. 2B). Treating T47D cells with estrogen or progesterone only induced phosphorylation of AKT, but not other signaling pathways (Fig. 2B). We screened a panel of inhibitors targeting JAK/STAT, AKT, or NF-κB and found that an inhibitor specific to inhibit Janus Kinases 1 and 2 (JAK1/2) was able to block STAT5 phosphorylation (Fig. 2C) and *Il2* expression (Fig. 2D). We also performed siRNA-based silencing of STAT5 (KD), which resulted in 60% reduction of STAT5 (sFig. 2E). STAT5 KD led to a substantial reduction in *Il2* expression in both vehicle- and PRL-treated groups (Fig. 2E). These results underscore the pivotal involvement of the PRL-induced STAT5 activation in *Il2* expression within MECs.

We compared *Il2*+ versus *Il2*-LMECs or BMECs using our scRNA-seq data and identified 28 common upstream regulators enriched in the *Il2*+ MECs (Fig. 2F), among which the prolactin was one of the top upstream regulators that are potentially involved in *Il-2* expression (Fig. 2G). Other common upstream regulators include interferon signaling (type I and type II IFN, as well as JAK1/2 and STAT pathways), T cell receptor signaling (CD3), inflammatory signaling (NFKB1A, IL1B, TNFSF10) as listed in Fig. 2G.

### Epithelial IL-2 modulates immune homeostasis within the mammary glands

To investigate the function of epithelial IL-2, we crossed *Il2*^fl/fl^ mice with *MMTV-Cre* to produce *Il2*^fl/f^/MMTV-Cre progeny (MEC-cKO). IL-2 protein expression was determined in LMECs or BMECs from WT (*Il2*^fl/fl^) and MEC-cKO mammary glands and the successful depletion of IL-2 was confirmed in both LMECs and BMECs from MEC-cKO mice relative to WT mice (Fig.3A-3B). T cells from the mammary glands, however, expressed equal amount of IL-2 protein (Fig.3A-3B), supporting specific deletion of IL-2 in MECs. A thorough datamining of scRNA-seq from the human single-cell RNAseq dataset failed to detect IL-2 receptors on the MECs (sFig. 3A), suggesting that epithelial IL-2 works on other cell types – mostly T, NK or other lymphocytes. We performed a thorough immune profiling of mammary glands from WT or MEC-cKO virgin female mice (sFig. 3B, gating strategy for lymphocytes in the mammary glands). One of the most consistent changes is the substantial reduction of NK cells (Fig. 3C-3E, NK cells – CD11b^-^ CD3^-^ CD19^-^ NK1.1^+^ in the lymphocyte gating), which is restricted to the mammary glands because NK cell number in other organs and tissues (spleen, lymph nodes, and blood) were not changed (Fig. 3F). We checked IL-2R expression levels on NK cells, and found that there are higher levels of IL-2RB and IL-2RG in human NK cells through human protein atlas(sFig. 3A) and mouse NK cells using flow cytometry (sFig. 4A). IHC staining for NK 1.1 revealed that NK cells were situated on the BMEC layer, in close proximity to LEMCs (sFig 4B). Moreover, we further confirmed that the NK1.1^+^ population in the mammary gland is predominantly NK cells rather than ILC1. This conclusion is supported by the majority of these cells being CD49a^-^CD127^-^ (markers indicative of ILC1), and the identification of two NK cell subpopulations characterized by relative expression levels: CD49b^high^EOMES^mid^ and CD49b^low^EOMES^high^ (sFig. 3B). The NK cells from mammary glands of MEC-cKO mice also exhibited an active phenotype as evidenced by the intracellular staining of IFN-γ, but the IFN-γ production by T_h_1 CD4conv and CD8^+^ T cells was not influenced by MEC-cKO of IL-2 (Fig. 3G-3I). To our surprise, MEC-cKO of IL-2 did not lead to significant changes in T cell populations including regulatory T cells (Tregs, CD4^+^CD25^+^FOXP3+) (sFig. 5A-5B), non-Treg CD4+ T cells (sFig. 5C-5D, CD4conv – CD4^+^FOXP3^-^), CD8^+^ effector memory T cells (sFig. 5E-5F, CD8 Tem – CD8^+^CD44^+^CD62L^-^), and B cells (sFig. 5G-5H, CD19^+^B220^+^).

**Figure 3.**
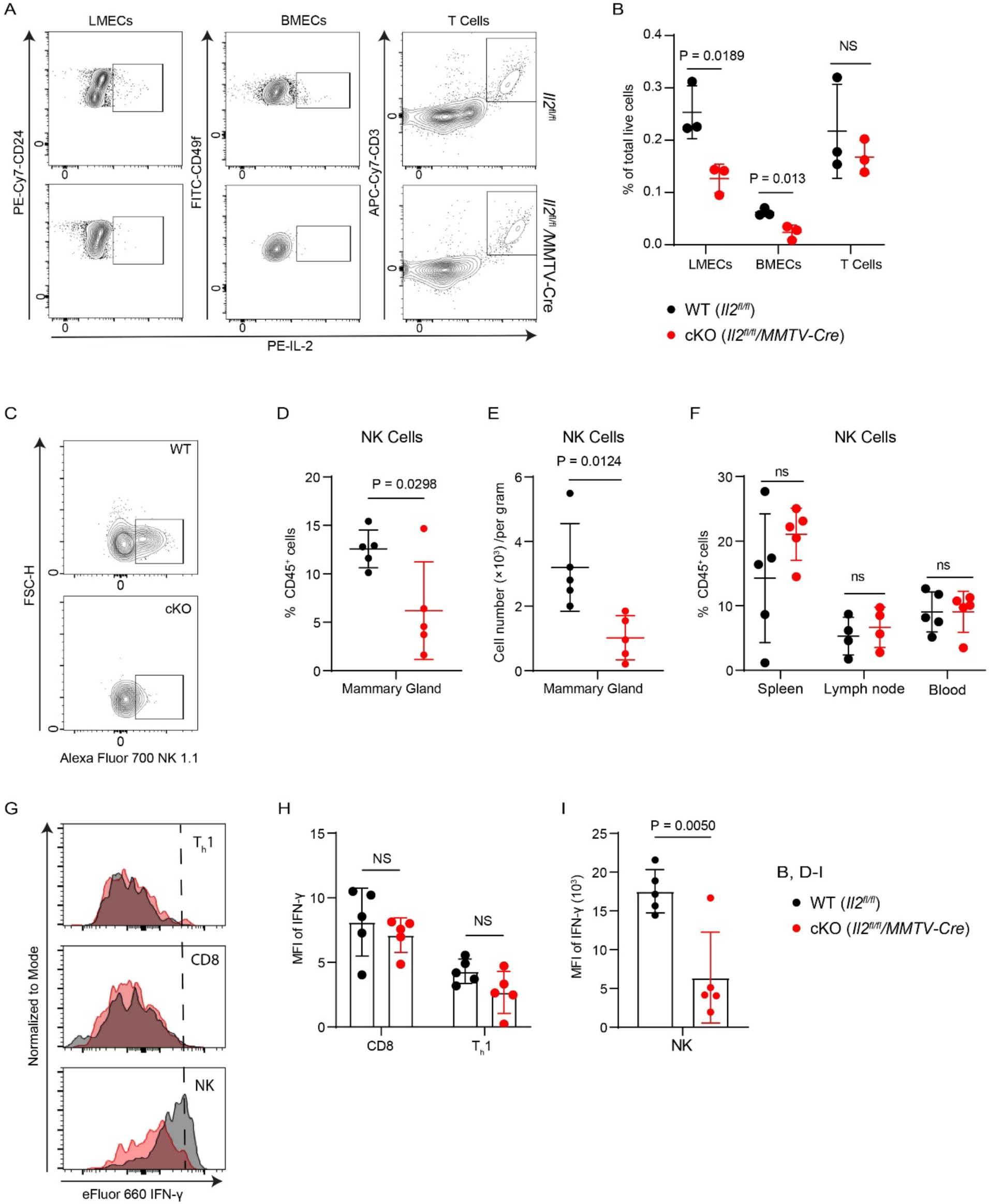
MEC-IL-2 depletion results in the downregulation of NK cell number and function. **A.** The schematics of IL-2 expression of LMECs, BMECs and T cells in *Il2*^fl/fl^ and *Il2*^fl/fl^/MMTV-Cre mice. **B.** IL-2 positive cell percentage of total live cells in mammary glands. **C**. The schematic of NK cells gating in the MG for flow cytometry analysis. **D.** NK cell percentage of CD45^+^ cells in the MG. **E.** NK cell number (per gram) in the MG. **F.** NK cell percentage of CD45^+^ cells in the spleen, lymph node, and blood. **G.** The histogram of IFN-γ in T_h_1, CD8, and NK cells for flow cytometry analysis. **H.** MFI (median fluorescence intense) of IFN-γ of T_h_1 and CD8 cells in the MG. **I.** MFI of IFN-γ of NK cells in the MG.

Active T cells are acknowledged as the primary source of IL-2. For comparison, we also produced *Il2 ^fl/fl^/Lck-Cre* (T-cKO) mice that have T cell-specific knockout of IL-2. We confirmed the IL-2 depletion from T cells (Fig. 4A-4B). We also observe a consistent reduction of NK cell numbers (Fig. 4C) and activation status evidenced by the intracellular staining of IFN-γ (Fig. 4D-4E). Unlike MEC-specific IL-2 depletion, T-cKO of IL-2 systematically impaired NK cell proliferation, significantly reducing NK cell numbers in the spleen, lymph nodes, and blood (Figure 4F). Additionally, T-cKO of IL-2 resulted in significant changes of several other lymphocytes including a significant decrease in Tregs (sFig. 6A-6B), a significant increase in CD4conv cells (sFig. 6C-6D) and in CD8 Tem cells (sFig. 6E-6F), as well as a substantial reduction in B cells (sFig. 6G-6H, CD19^+^B220^+^), consistent with previous publications.^13, 31^

**Figure 4.**
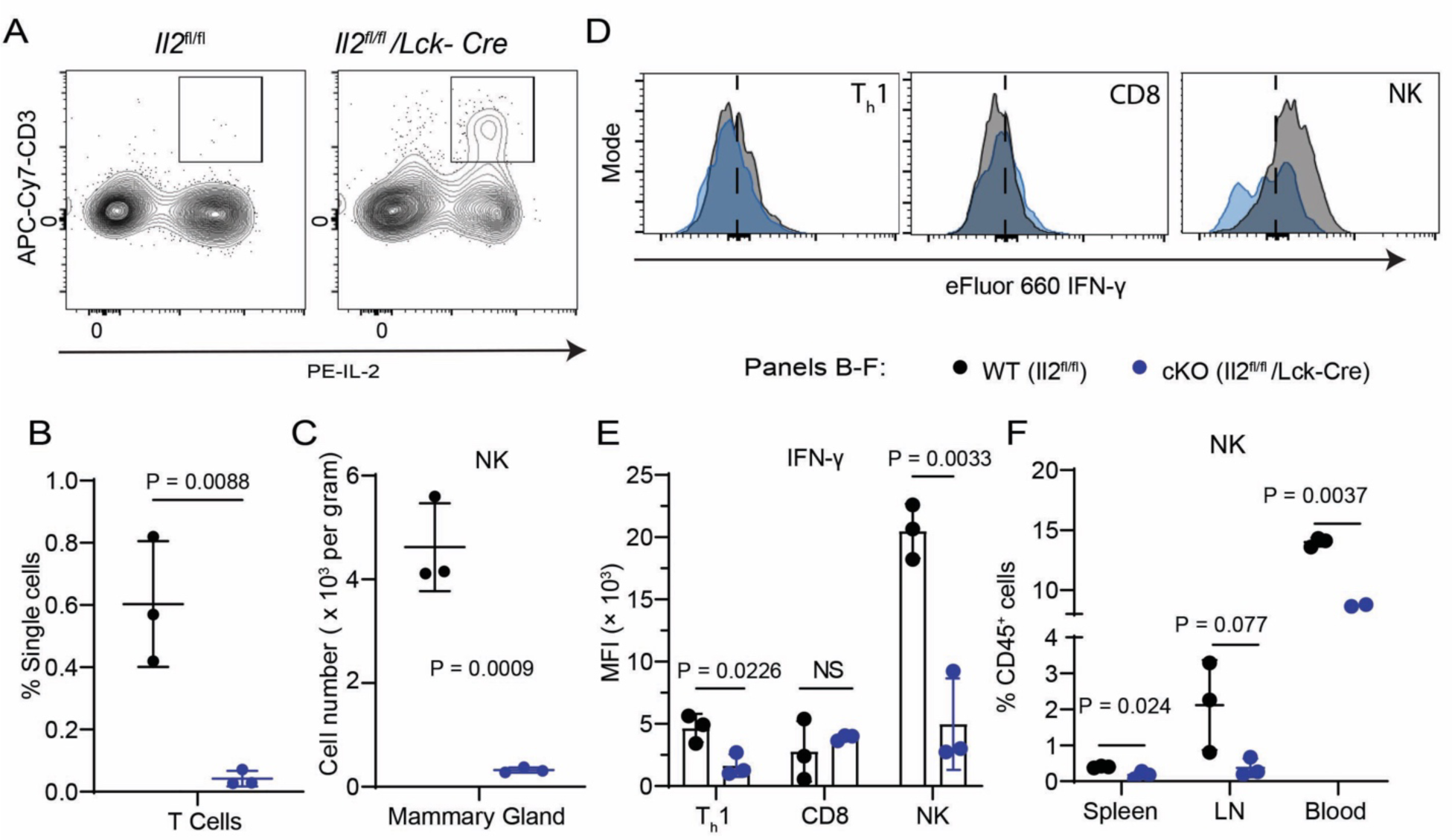
T cell-IL-2 depletion results in global reduction of NK cell number. **A.** The schematic of T cells gating between WT and T-cKO mice for flow cytometry analysis in the spleen. **B**. IL-2 positive T cell percentage of single cells between WT and T-cKO mice in the spleen. **C.** NK cell number (per gram) in the MG between WT and T-cKO mice. **D.** The histogram of IFN-γ in T_h_1, CD8, and NK cells for flow cytometry analysis between WT and T-cKO mice. **E.** MFI of IFN-γ of NK cells in the MG between WT and T-cKO mice. **F.** NK cell percentage of CD45^+^ cells in the spleen, lymph node, and blood between WT and T-cKO mice.

It has been shown that MMTV-Cre transgene can be expressed in other immune cells including lymphocytes. We performed a side-by-side comparison of Cre expression driven by Lck-Cre or MMTV-Cre transgene using flow cytometry of peripheral blood. Lck-Cre specifically drove Cre expression in T cells, but not in other cell types such as B cells, NK cells, or DCs (sFig. 7A-B). MMTV-Cre, on the other hand, did not drive any significant expression of Cre in any of these immune cell types in the blood (sFig. 7A-B), supporting that MMTV-Cre drives Cre expression mainly in the mammary epithelial cells. Our data agrees with published data that showed the leakage of MMTV-Cre only occurs in a small percent of common lymphoid progenitors (CLPs, less than 5%) using a lineage tracing experiment ^32^.

These data support that MEC-IL-2 – together with T cell IL-2, plays a significant role in the NK cell function within the mammary glands; T cell-produced IL-2, however, plays a much broader role in lymphocyte regulation within the mammary glands.

### Epithelial IL-2 is critical for immunosurveillance within mammary glands

Our results identified a significant loss of NK cells within the mammary glands of the MEC-cKO mice. NK cells are vital innate lymphocytes involved in cancer immunosurveillance via cytotoxic functions. There is no direct evidence that NK cells are involved in the pathophysiology of mammary glands, but the IFN-γ/STAT1 signaling pathway – one of the major effector pathways of NK cells – is known as tumor suppressive pathway because genetic deletion of STAT1 leads to estrogen receptor-positive luminal breast cancer ^33^ and other cancers ^34^. We confirmed the significant *IFNGR1* and *IFNGR2* expressions in LMECs and BMECs in our scRNA-seq dataset (sFig. 8), supporting that MECs are able to receive IFN-γ for STAT1 activation. Previous proteomic data also revealed IFNγR1 expression exclusively in luminal progenitors in vivo^35^. We examined the potential impact of MEC-cKO of IL-2 on MECs under physiological conditions. MEC-cKO of IL-2 led to an imbalance of LMEC populations relative to BMEC cells, with a marked increase of LMECs in the cKO mammary glands (Fig. 5A-5B). In agreement with the literature, the expansion of LMECs is likely due to the decreased level of the STAT1 activation in the MEC-cKO mouse – as evidenced by the alteration in tyrosine phosphorylation of STAT1 in MECs (Fig. 5C), supporting that IFN-γ primarily activates STAT1 in the luminal lineage. It has been shown that earlier hyperplastic expansion of LMECs can be the cell-of-origins for breast cancers ^36–38^. Gross tissue examination of mammary glands from the 14-week-old MEC-cKO mice also confirmed the aberrant expansion of side branches within the ductal tree relative to a smoother look of those in WT females (Fig. 5D-5E).

**Figure 5.**
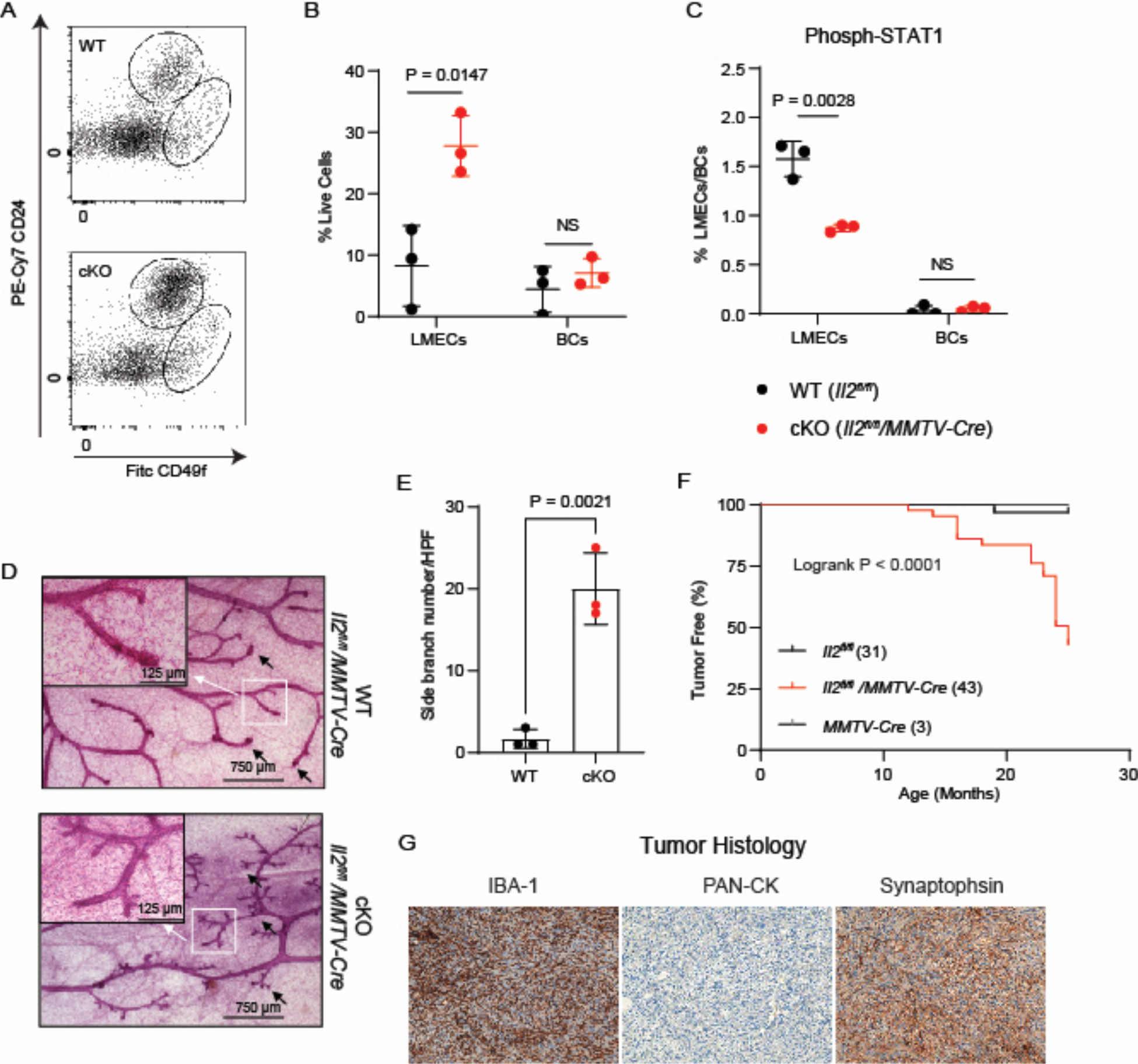
Loss of tumor immunosurveillance in the aging MEC-cKO mice. **A.** The schematic of MEC gating for flow cytometry analysis. **B.** Percent of BMEC or LMEC in the mammary glands. **C.** Percent of STAT1 active epithelial cells as indicated by tyrosine phosphorylation of STAT1. **D.** Images of whole mount staining of WT and cKO mammary glands (bar = 750 µM in low magnification and bar = 125 µM in the left top panels). **E.** Quantification of side branch number from the WT or cKO mammary glands. **F-G.** Spontaneous tumor development in the aging cKO mice. **(F)** Survival curve of total tumor free mice among *Il2^fl/fl^* (WT), *MMTV Cre* (WT), and *Il2^fl/fl^ MMTV Cre* (cKO); and (**G)** Representative tumor IHC staining with ionized calcium-binding adapter molecule 1 (Iba-1), pan-cytokeratin (PAN-CK), and synaptophysin. B,C and E: two-sided unpaired *t* test was performed, with P values indicated. NS: not significant. F. Logrank test for Kaplan-Meier survival was used.

To investigate the long-term effect of MEC-cKO of IL-2 on mammary tumorigenesis due to the potential loss of immunosurveillance from NK cells, we monitored both WT and MEC-cKO mice for up to two years of age. Notably, cKO mice began to develop solid tumors at mammary glands and/or other sites as they aged, with the earliest case being observed in a mouse of 12 months old (Fig. 5F). The control group had only one mouse with a tumor (Fig. 5F). IHC staining of tumors showed synaptophysin positivity, ionized calcium-binding adapter molecule 1 (Iba-1) positivity, and pan-cytokeratin negativity, supporting that the tumor is none-epithelial origin and likely with neuroendocrine features (Fig. 5G). The timeline of tumor development in the MEC-cKO group was consistent with that of the STAT1-deficient mice^33^.

To explore the relationship between tumorigenesis and alterations in the immune system in MEC-cKO mice, we examined the immune phenotypes of the aged mice (22 months old). We observed a consistent loss of NK cells in the mammary glands of MEC-cKO mice (Fig. 6A-6B), with an unexpected reduction of NK cells in the peripheral blood but not in the spleen (Fig. 6C). IFN-γ production was also reduced from the NK cells in MEC-cKO mice (Fig. 6D-6E). The reduction of NK cells in the blood may explain why tumors developed from various tissues other than the mammary glands.

**Figure 6.**
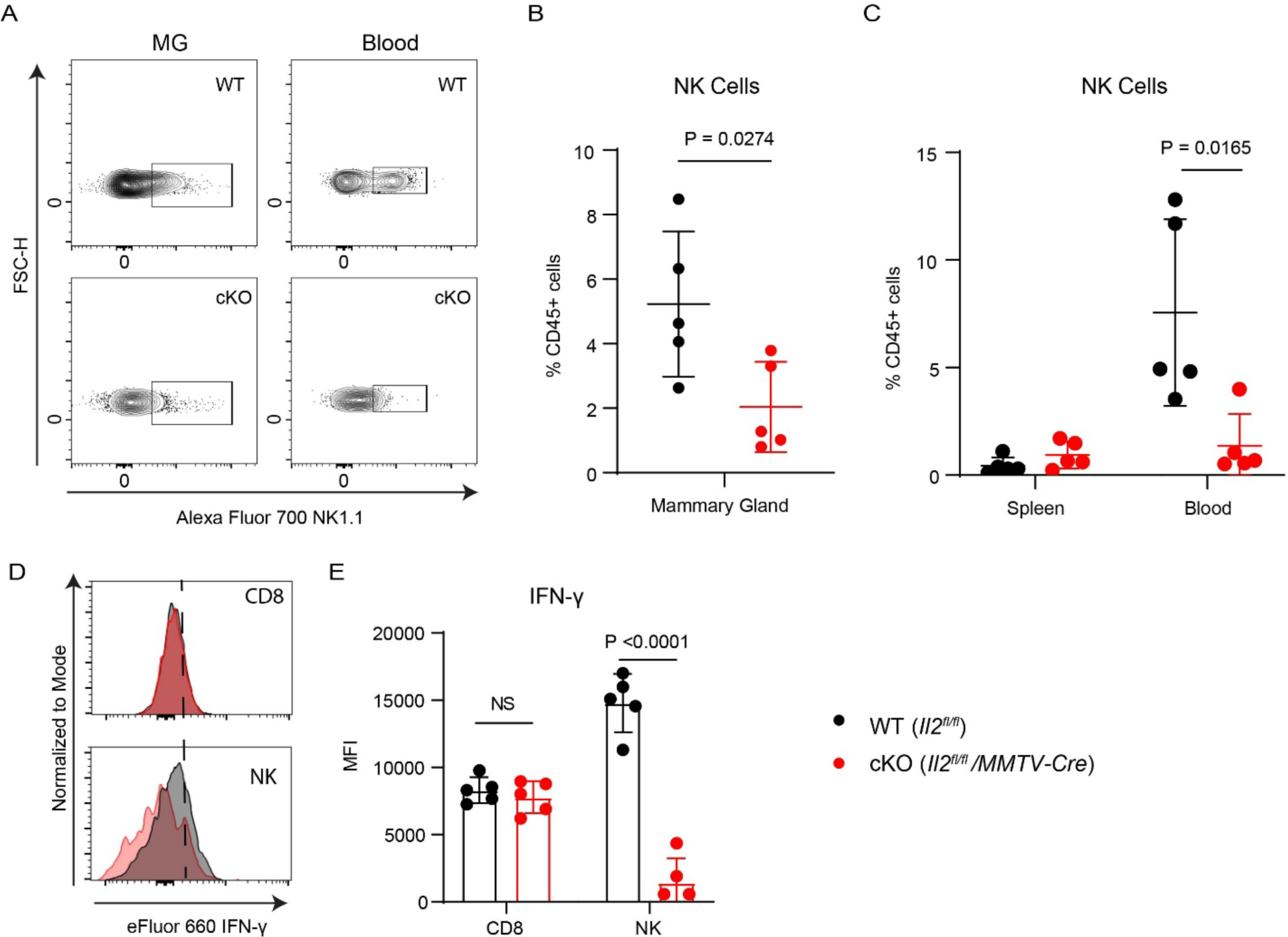
NK cells are significantly impaired in the aging MEC-cKO mice. **A.** The schematic of NK cells gating in the MG and blood from the aging mice (20-month-old) for flow cytometry analysis. **B.** Percent of NK cells within the total CD45^+^ cells in the MG from aging mice. **C.** Percent of NK cells within the total CD45^+^ cells in the spleens and blood from the aging mice. **D.** The histogram of IFN-γ in CD8, and NK cells from aging mice for flow cytometry analysis. **E.** MFI of IFN-γ of NK cells in the MG from aging mice. The two-sided unpaired *t* test was performed, with P values indicated. NS, not significant.

## Discussion

In our study, we made a groundbreaking discovery that MECs represent a novel source of IL-2. This highlights the capability of MECs to maintain immune homeostasis within the MGs via producing IL-2. MG consists of a complex structure, comprising inner luminal and outer basal layers, embedded within a fat pad housing adipocytes, fibroblasts, and immune cells^39–41^.

Functioning as an endocrine organ, MG undergoes dynamic changes, particularly during pregnancy, lactation, and involution. Hormones such as estrogen, progesterone, and prolactin regulate this process, necessitating the maintenance of immunotolerance and immunosurveillance against self-antigens and the commensal or pathological microbes^42–46^. MEC-produced IL-2 is critical in maintaining the immune homeostasis of the MG.

We initially hypothesized that hormones play a pivotal role in MEC-mediated IL-2 expression, due to the impact of female hormones on MG physiology and function. Because we detected IL-2 expression across both LMECs and BMECs, we believe that estrogen and progesterone should not be considered as the primary instigators of MEC-IL-2 expression, given the exclusive localization of their receptors (ER and PR) within LMECs, excluding BMECs^47^. Prolactin, which interacts with PRLR expressed across various cell types, emerges as a key regulator of MEC-IL-2 expression, validated in T47D and HC11 cell line models. Prolactin’s interaction with PRLR leads to STAT5 phosphorylation, regulating gene expression in MECs to support milk synthesis, MG development, and component secretion^48–50^. The phosphorylation of STAT5 is a critical factor in the positive loop of T cell activation. Elevated IL-2 can stimulate T cells through IL-2R binding, triggering STAT5 phosphorylation and converting CD4 and CD8 T cells into short-lived effector T cells that simultaneously express IL-2. This underscores the importance of the prolactin-STAT5 axis in MEC-IL-2 expression^3, 51^.

MMTV-Cre (line D) is the most commonly used Cre line for deletion of target floxed alleles in the mammary epithelial cells, whose expression has been shown to be mainly restricted to mammary epithelium and salivary epithelium^52^. MMTV-LTR-driven gene expression can occur in several other tissues at low frequency from earlier studies^52–54^, but a careful lineage tracing experiment showed that MMTV-Cre expression occurs in less than 5% of common lymphoid progenitors^32^, indicating that MMTV-Cre-induced IL-2 deletion only occurs in very low percent of lymphocytes. Our study further validated published observations and showed no difference in IL-2^+^ T cells upon activation comparing T cells coming from *Il2*^fl/fl^ and MMTV-Cre/*Il2*^fl/fl^ mice. The lack of Cre expression in the MMTV-Cre mice within different lineages (sFig. 7) further excludes the significant impact of IL-2 deletion from other cell types on NK cells within the MG. We did not detect any significant change in T cell subpopulations, further excluding the involvement of T cell-produced IL-2 in the immunological changes within the MG.

As primary consumers of IL-2, Treg cells predominantly express the high-affinity IL-2 alpha receptor (CD25/IL2RA)^55^. Intriguingly, while T-cKO of IL-2 disrupts immune homeostasis, MEC-cKO of IL-2 did not significantly affect the proliferation of Treg, CD8^+^ Tem, CD4conv, and B cells. This suggests that T cell-IL-2 can compensate for the absence of MEC-IL-2 on these immune cells; whereas NK cells are the primary effector cell population of MEC-IL-2.

Within the MG, NK cells are highly responsive to changes in IL-2 expression levels. NK cells express the highest-level IL-2RB (CD122) and IL-2RG (CD132) (sFig. 3A and sFig. 4A), resulting in the hypersensitivity of NK cells to the MEC-IL-2 alteration. Presumably, the high level of IL-2RB and IL-2RG are sufficient to induce IL-2 dependent signaling transduction – with or without IL-2RA – as reported before^56^. IL-2RB knockout exhibits a major loss of NK cells, but effector T cells can be spontaneously activated, supporting the critical role of high IL-2RB in NK cell function.^57^ Moreover, NK cells in the MG are positioned close to MECs, actively consuming and responding to MEC-derived IL-2 as the initial cell population over time. IL-2 stimulates NK cells to produce IFN-γ, establishing a crucial defense against tumor growth and showing a positive correlation with favorable cancer prognosis^58, 59^. In the MEC-IL-2 depletion model, we observed a reduction in NK cell frequency and IFN-γ production, correlating with tumor initiation. Furthermore, our findings, along with those of other studies, support the role of IFN-γ in restricting the expansion of LMECs and inhibiting mammary morphogenesis by limiting branching^22, 35^. In MEC-cKO mice, mammary ductal differentiation went awry, potentially increasing the risk of certain diseases, notably breast cancer^60^.

In conclusion, our research unveils a novel communication mechanism between MECs and NK cells. We have shown that the prolactin-STAT5 pathway induces MECs to produce IL-2, thereby supporting NK cell regiment. This discovery introduces a new source of IL-2 and adds a crucial component to the well-established mechanism governing NK cell immunosurveillance. Ultimately, our study provides fresh insights into cancer development, which is regulated by the crosstalk of local epithelial cells and immune cells.

## Methods

### Single-cell RNA analysis

We utilized multiple datasets from GEO Dataset: GSE143159^23^. We imported the filtered gene count matrix and performed quality control checks on all samples as part of the standard workflow used by the Seurat R package, version 2.3.4 in R. Any cells that exhibited unique RNA feature counts below 200 or a percentage of mitochondrial DNA exceeding 5% were excluded from the analysis. scRNA-Seq clustering and visualization was described previously^23^.

### Cell lines and cell culture

Human cell lines: T47D (HTB-133, ATCC) were cultured in RPMI-1640 (R8758, Sigma) and supplemented with 0.2 unit/ml bovine insulin (I0516-5ML, Sigma), 10% Fetal Bovine Serum (FBS, F2442, Sigma) and 100 U/ml penicillin and 100 µg/ml streptomycin (P4333, Sigma).

Mouse cell lines: HC11 cells (CRL-3062, ATCC) were cultured in RPMI-1640 and supplemented with 10% FBS (F2442, Sigma) and 100 U/ml penicillin, and 100 µg/ml streptomycin (P4333, Sigma).

HC11 cell line were differentiated based on an established protocol^30^. 400,000 HC11 cells per well of a 6-well plate were seeded. When the cells reach approximately 90-100% confluence, the medium was replaced with a medium containing FBS and insulin for 24 hours, following with the replacement of differentiation medium (HIP medium, RPMI-1640 supplemented with 10% FBS, 1 μg/mL Hydrocortisone, 5 μg/mL Insulin, and 5 μg/mL Prolactin) for up to 10 days. Medium were changed every 2-3 days for HIP-treated cells and control cells. The formation of “domes”-like growth pattern indicated successful differentiation. To detect IL-2 expression using flow cytometry, an activation cocktail with Brefeldin A (4233043, 1:500, Biolegend) was added for 3 hours before harvesting cells.

Ruxolitinib (941685-37-6, Sigma), a JAK1/2 inhibitor, was used to inhibit JAK1/2 tyrosine kinases. Briefly, T47D cells were pretreated with 0, 1, and 5 μM of ruxolitinib for 48 hrs, following with 10 nM of PRL treatment. Western Blotting analysis was employed to assess the STAT5 and phospho-STAT5 using specific antibodies and qPCR to measure the expression of *IL2* mRNA.

### STAT5 knockdown (KD) assay

T47D cells were transfected with 25 nM of ON-TARGETplus Human STAT5A siRNA (L-005169-00-0005, Horizon) or ON-TARGETplus Non-targeting Control Pool (D-001810-10-05, Horizon) using GeneTran III reagent (GT2211, Biomiga). After 48 hours, we replaced the medium and cultured the cells for an additional 24 hours to allow for efficient KD.

### Western Blotting

Cell lysates were collected using radioimmunoprecipitation assay buffer (RIPA) lysis buffer (150 mM NaCl, 5 mM EDTA, 50 mM Tris pH 8.0, 0.5% sodium deoxycholate, 1% NP-40, 0.1% SDS) supplemented with the protease-phosphatase inhibitor (PPC1010, Sigma). The lysates were separated by SDS-PAGE and analyzed using standard Western Blotting procedure^61^. Antibodies: STAT-5 (94205S, Cell Signaling, 1: 1000), phos-STAT5 (9359S, Cell Signaling, 1: 1000), NF-κB (8242S, Cell Signaling 1:1000), phos-NF-κB (3303S, Cell Signaling, 1: 1000), AKT (9272S, Cell Signaling, 1:1000), phospho-AKT (4060S, Cell Signaling, 1: 1000), ERK1/2 (4695S, Cell Signaling, 1: 1000), phospho-ERK1/2 (4370S, Cell Signaling, 1: 1000), GAPDH (5174S, Cell Signaling, 1: 5000).

### Quantitative PCR (qPCR)

RNA was extracted from cells using the RNeasy Mini Kit (QIAGEN, cat. no. 74104). Subsequently, the mRNA was converted into cDNA as template for qPCR, using the random hexamers method (ThermoFisher). To assess the relative expression levels of specific genes, human *IL2* (PN4351370, FAM, ID: Hs00174114_m1), human *GAPDH* (PN4448485, VIC, ID: Hs02786624_g1) TaqMan primers, TaqMan Fast Advanced Master Mix (4444557, Thermo Fisher) and the ΔΔCt method of qPCR were used. The amplification of *GAPDH* serving as an endogenous control for normalization.

### Animal experiments

All animal studies were approved by the University of Florida Institutional Animal Care and Use Committee (IACUC) under protocol 202110399. All animals were housed in Pathogen-Free vivarium of an Association for Assessment and Accreditation of Laboratory Animal Care accredited facility at the University of Florida and performed in accordance with IACUC guidelines. The room temperature is 21.1–23.3 °C with an acceptable daily fluctuation of 2 °C. Typically the room is 22.2 °C all the time. The humidity set point is 50% but can vary ±15% daily depending on the weather. The photoperiod is 12:12 and the light intensity range is 15.6-25.8 °C.

C57BL/6 *IL2^f/f^* mice were initially made in Dr. Smith group^31^. *Tg(MMTV-cre)4Mam/J* ^52^ and *B6.Cg-Tg(Lck-cre)548Jxm/J* ^62^ were purchased from Jackson Laboratories (Bar Harbor, ME).

Mice were euthanized in accordance with IACUC protocol and tissues were collected for further analysis.

### Mouse mammary tissue dissociation

#4 inguinal mammary glands from mice were used after removal of the inguinal lymph node, carefully excised using a scalpel and digested with gentle collagenase/hyaluronidase (07919, STEMCELL). To maximize viable cell yield, we determined the optimal enzyme concentrations and digestion times. For immune cells, we diluted one part gentle collagenase/hyaluronidase with four parts of complete EpiCult^TM^-B medium (05610, STEMCELL) supplemented with 5% FBS to digest for two hours. For MECs, we diluted one part gentle collagenase/hyaluronidase with nine parts of complete EpiCult^TM^-B medium to treat for 8 hours. Following digestion, tissues were centrifuged at 350 × *g* for 5 minutes. The cell pellet was resuspended in a mixture of one part cold HBSS (37150, STEMCELL) supplemented with 2% FBS (HF) and four parts of Ammonium Chloride Solution (composed of 8.3 g NH_4_Cl, 1 g KHCO_3_, 1.8 mL of 5% EDTA, and ddH_2_0 to make 1 L), centrifuged at 350 × *g* for 5 minutes. Tissue pellets were washed again with cold HF and centrifuged, further digested with pre-warmed Trypsin-EDTA and gently mixed it by pipetting for 1 - 3 minutes. Cells were pelleted and resuspended in 2 mL of pre-warmed dispase (5 U/mL; 07913, STEMCELL) and 200 µL of DNase I Solution (1 mg/mL; 07900, STEMCELL), digested for 1 minute to further dissociate into single cells. Cells were then centrifuged at 350 × *g* for 5 minutes, resuspended with 10 ml of cold HF and filtered it through a 40 µm Cell Strainer (27305). Cells were centrifuged at 350 x *g* for 5 minutes and ready for use in subsequent experiments.

### Whole mount of the mammary glands

The whole mount staining was based on an established protocol^63^ with slight modifications. Briefly, staining solution was prepared by dissolving 1 g of carmine and 2.5 g of aluminum potassium sulfate in 500 ml of distilled water, then boiling for 20 minutes in a 1-liter Erlenmeyer flask. The final volume was adjusted to 500 ml with water and filtered through Whatman paper with a small amount of thymol as a preservative. Mouse mammary glands were excised and carefully spread on a glass slide, and emerged into the Carnoy’s fixative (composed of 100% ethanol, chloroform, and glacial acetic acid in a 6:3:1 ratio) for 4 hours at room temperature, or alternatively at 4°C overnight. After fixation, slides were washed with 70% ethanol for 15 minutes and then gradually rehydrated using 70%, 35%, and 15% ethanol and water baths for 5 minutes each. After final rinse with distilled water for 5 minutes, slides were stained in the carmine alum solution overnight at room temperature. Following staining, the slides were gradually dehydrated through sequential ethanol baths (50%, 70%, 95%, 100%) for 5 minutes each, cleared in Histo-clear overnight. Mammary glands were immersed in methyl salicylate and stored in a fume hood for imaging.

### Flow cytometry

Single-cell suspension was obtained from cultured cells or digested mammary glands. Cells were washed and incubated with combinations of the following antibodies: Fixable Viability Dye eFluor 780 (65086514, eBioscience), anti-mouse CD62L-BV785 (clone MEL-14), anti-mouse/human CD44-PerCP (clone IM7), anti-mouse CD11B-PEdazzle 594 (clone M1/70, 1:200), anti-mouse CD45-AF532 (clone 30F.11), anti-mouse CD3-APC/Cy7(clone 17A2), anti-mouse CD8-BV510 (clone 53-6.7), anti-mouse CD4-BV605 (clone GK1.5), anti-mouse NK1.1-AF700 (clone PK136), anti-mouse/human CD45R/B220-BV 570 (clone RA3-6B2), anti-mouse CD19-Alexa Fluor488 (clone 6D5), anti-mouse CD25-PE-Cy5 (clone PC61)), anti-mouse CD31-APC (W18222B), anti-mouse TER119-APC (TER119), anti-mouse CD24-PE/Cy7 (30-F1), anti-mouse CD49f-FITC (GoH3) plus zombie red (cell viability, Thermo Fisher) and mouse FcR blocker (anti-mouse CD16/CD32, clone 2.4G2, BD Biosciences), maintaining cell activation cocktail. After surface staining, cells were fixed and permeabilized using the FOXP3/Transcription Factor Staining Buffer Set (eBioscience). Cells were stained with a combination of the following antibodies: anti-mouse FOXP3-efluor 450 (clone FJK-16S, 1:50, eBioscience), anti-mouse/human T-bet-BV421 (clone 4B10), anti-mouse IL-2-PE (clone JES6-5H4). Flow cytometry was performed on a 3 laser Cytek Aurora Cytometer (Cytek Biosciences, Fremont, CA) and analyzed using FlowJo software (BD Biosciences). All antibodies are from Biolegend, unless otherwise specified. Most antibodies were used at 1:100 dilution for flow cytometry, unless otherwise specified.

### Statistics

Graphs and statistical analyses were performed using GraphPad Prism 9.4.0 unless otherwise specified. For comparisons involving three or more groups, a one-way ANOVA was conducted, followed by Dunnett’s multiple comparison test for specific group comparisons. Unpaired T tests were used to compare means between two groups.

## Data Availability

We have used several published datasets including our own dataset GSE143159^23^, scRNAseq data from the human protein atlas (https://www.proteinatlas.org/), Mouse ENCODE transcriptome data (BioProject: PRJNA66167). All other data and materials are available upon request.

## Author Contributions

Funding: W.Z.; Conception and design: W.Z., L.W., N.B., R.H.K., K.A.S.; Development of methodology: L.W., N.B., W.Z.; Analysis and interpretation of data: L.W., N.B., R.P.M., J.M., T.T., M.C.K., M.E.C., A.P.M., K.C., R.H.K., W.Z.; Writing, review, and/or revision of the manuscript: L.W. and W.Z. for first draft; W.Z., L.W., K.A.S., for rewriting, revision and editing. Supervision: W.Z.

## Acknowledgements

The work is partially supported by NIH grants CA200673 (W.Z.), CA203834 (W.Z.), CA260239 (W.Z.). W.Z. was also supported by an endowment fund from the Dr. and Mrs. James Robert Spenser Family and the Startup fund from the University of Florida Cancer Center; as well as the University of Florida Health Cancer Center Support Grant P30CA247796 for subsidy of core facilities.

## Supplementary Figure Legends

**Supplementary Figure 1.**
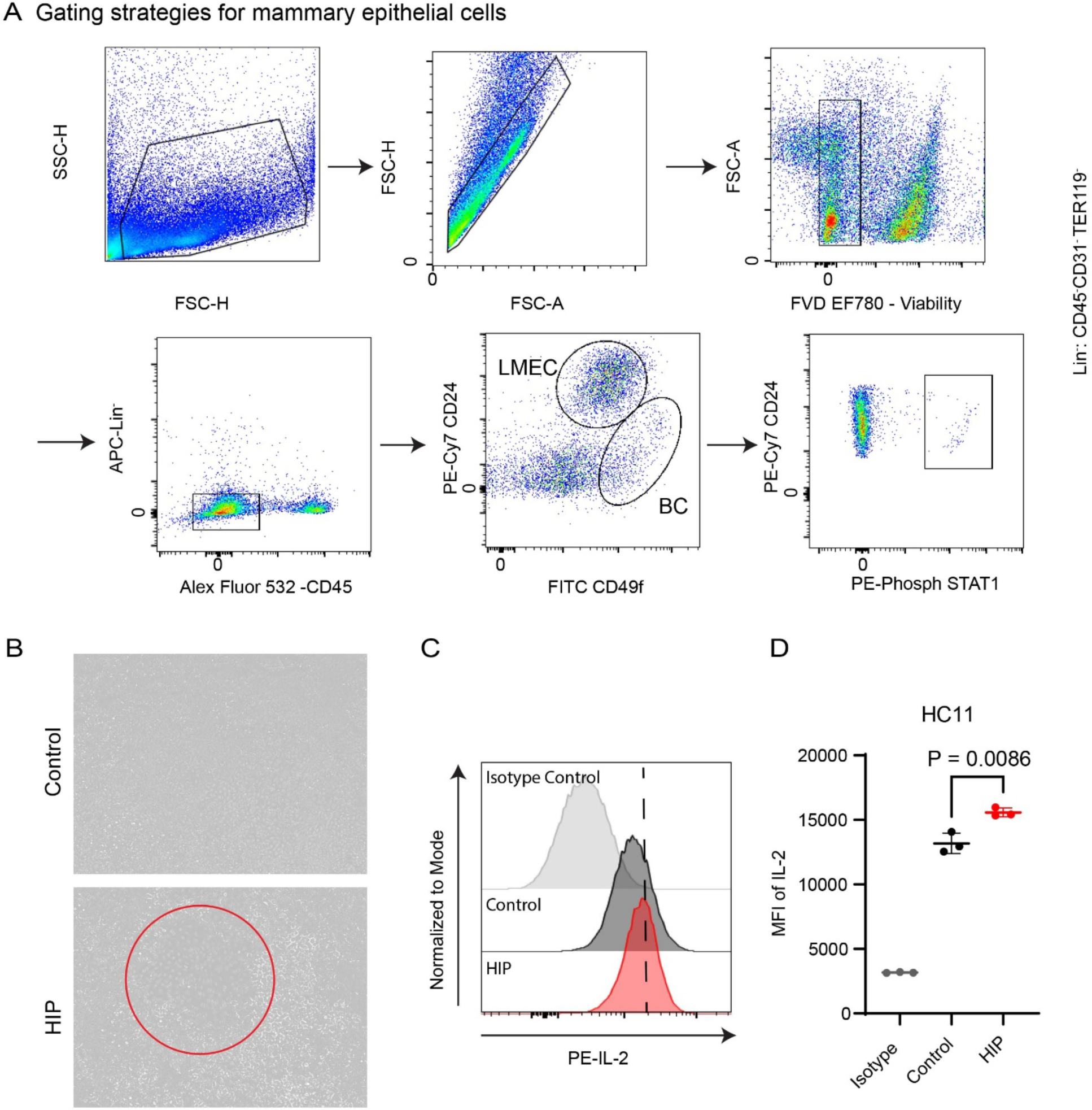
IL-2 expression in HC11, a mouse mammary epithelial cell line. **A.** Gating strategies for mammary epithelial cells. **B**. HC11 cell morphology between control and HIP medium after 10 days of incubation. **C-D.** IL-2 protein expression in mouse mammary epithelial cells. **(C)** representative flow cytometry of HC11 cells that were cultured in regular medium, or medium supplemented with hydrocortisone, insulin and prolactin (HIP). Isotype control, FMO + Isotype of HC11 cells cultured in HIP medium; Control, HC11 cells cultured in regular medium; HIP, HC11 cells cultured in HIP medium. **(D)** MFI of IL-2 summarized from **C** (n=3). Supplementary Figures for Figure 1 and Figure 2.

**Supplementary Figure 2.**
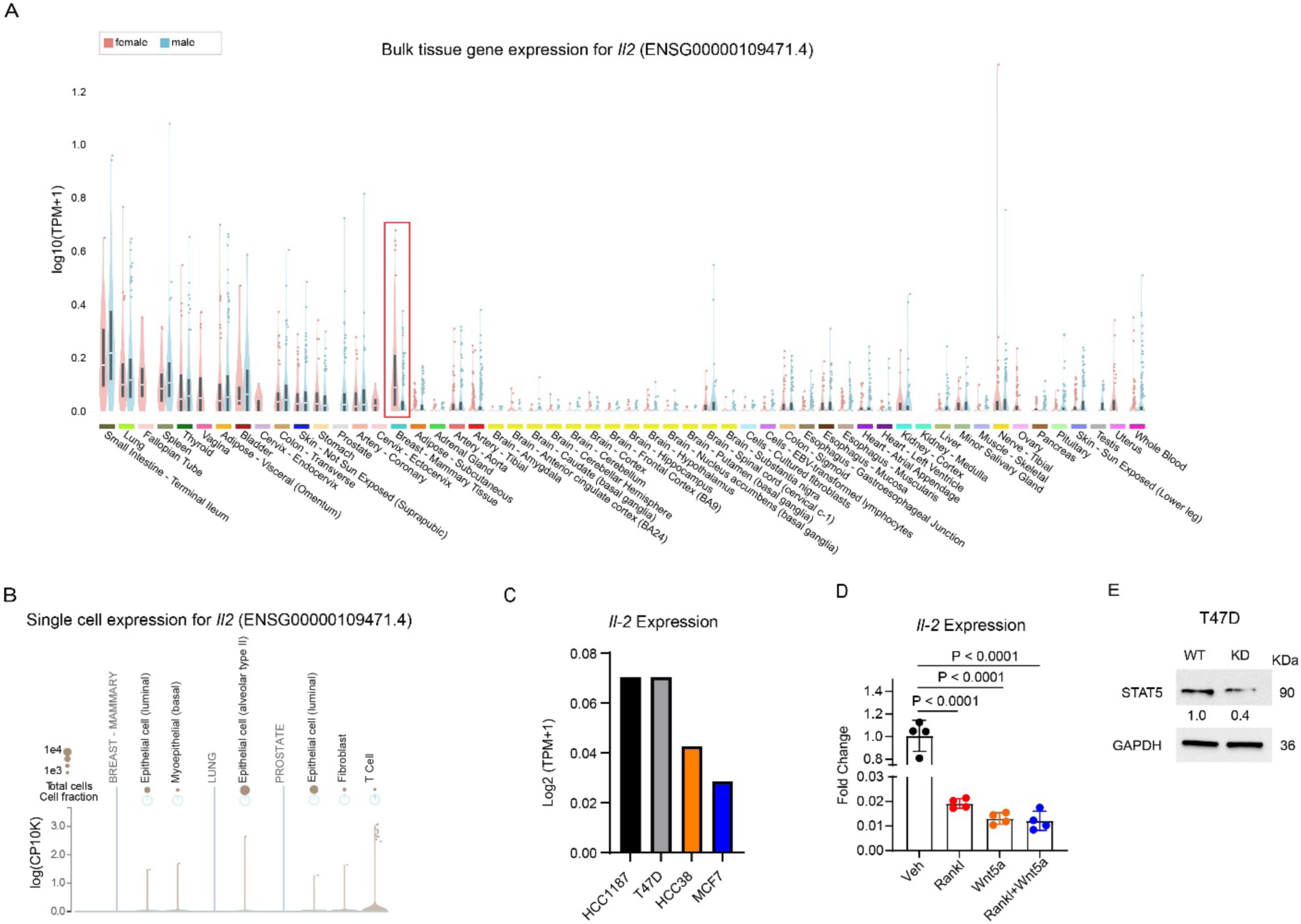
*Il2* mRNA expression is detected in human MECs. **A.** Bulk *IL2* mRNA expression in multiple tissues from man and woman (ENSG00000109471.4). **B.** *IL2* expression in breast, lung and prostate based on single cell RNAseq (ENSG00000109471.4). **C.** *IL2* mRNA expression in breast cancer cell lines, including HCC1187, T47D, HCC38 and MCF7 (Expression Public Access # 23Q4). **D.** RANKL and WNT5a inhibit *IL2* expression in the T47D cell line. **E.** STAT5 was knocked down by siRNA in the T47D cell line. Supplementary Figures for Figure 1 and Figure 2.

**Supplementary Figure 3.**
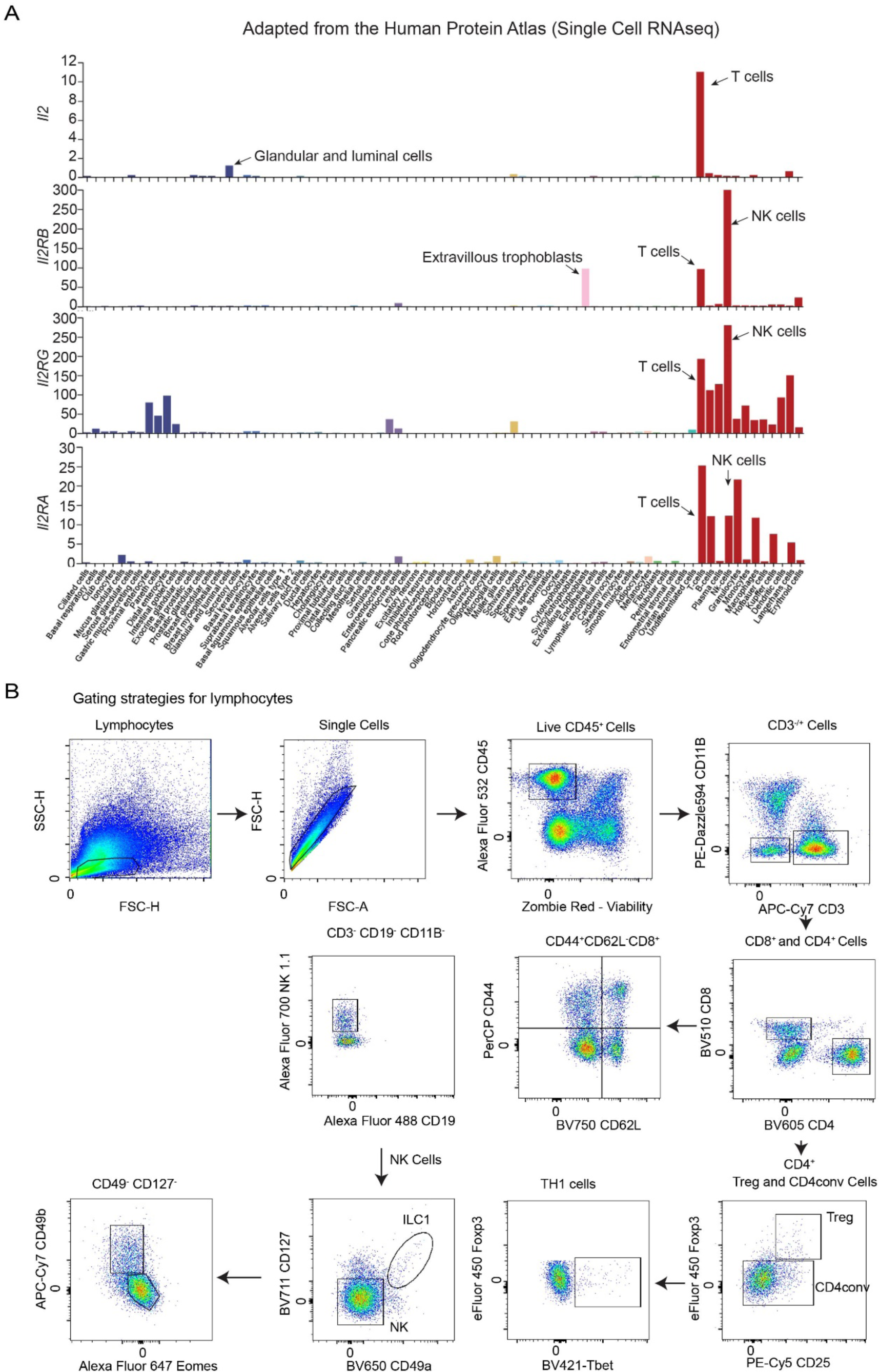
*IL2* and IL-2 receptors expression on different cell types from human tissues – adapted from the Human Protein Atlas**. A.** Single-cell RNAseq data from human protein Atlas showing the *Il2, Il2RB, Il2RG, and Il2RA* in different cell types. Note *IL2* mRNA is detected by glandular and luminal cells; whereas NK cells express the highest levels of *IL2RB* and *IL2RG*. **B.** Flow cytometry gating strategies for lymphocytes in the MG. Supplementary Figure for Figures 3-6.

**Supplementary Figure 4.**
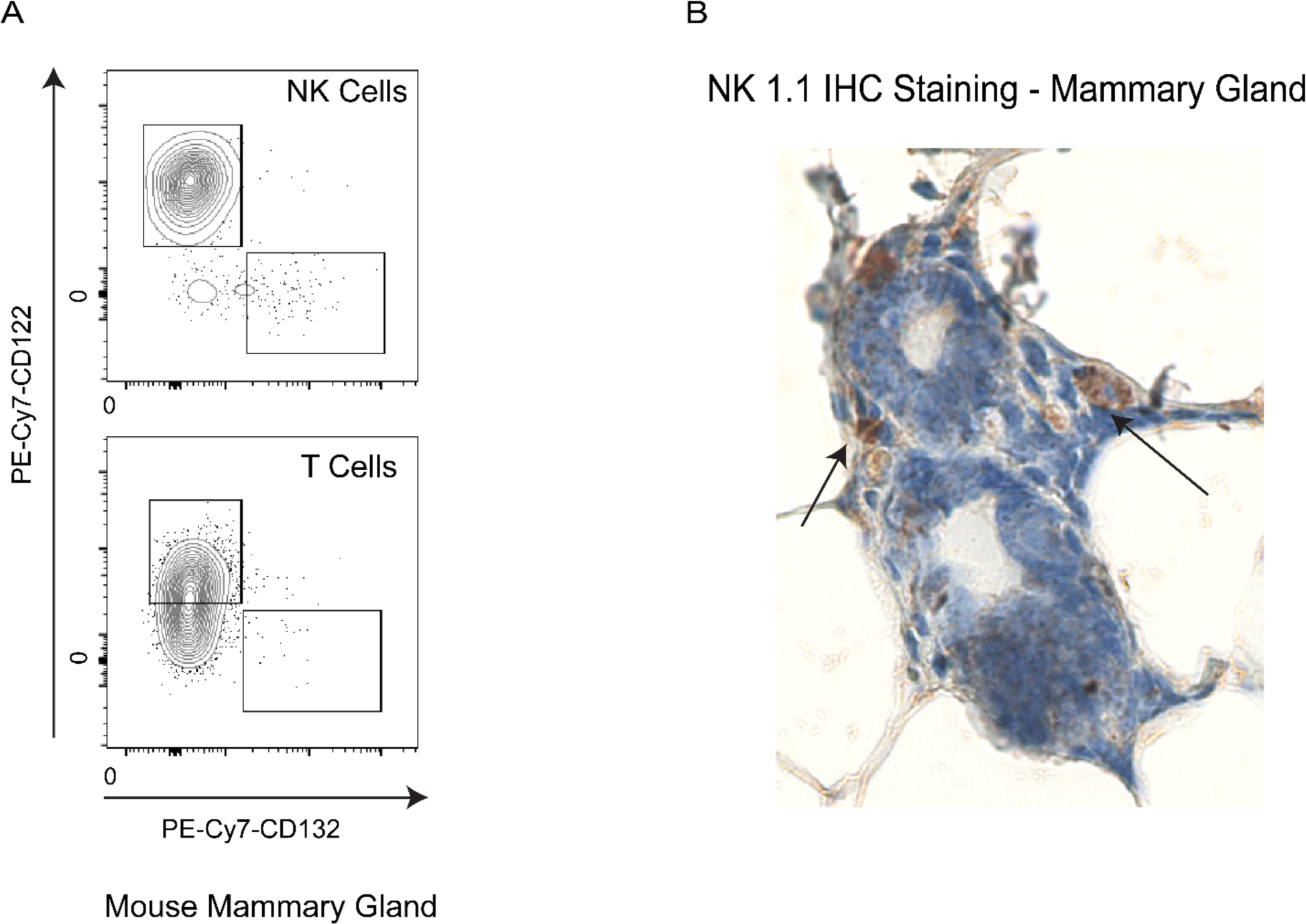
IL-2 receptors expression on NK cells and NK cells location in mammary ducts. **A.** CD122 (IL2RB) and CD132 (IL2RG) expression levels in NK cells and T cells from a representative mouse mammary gland. **B**. IHC stating of NK1.1 to identify the location of NK cells in mouse mammary ducts. Supplementary Figure for Figure 3 and 4.

**Supplementary Figure 5.**
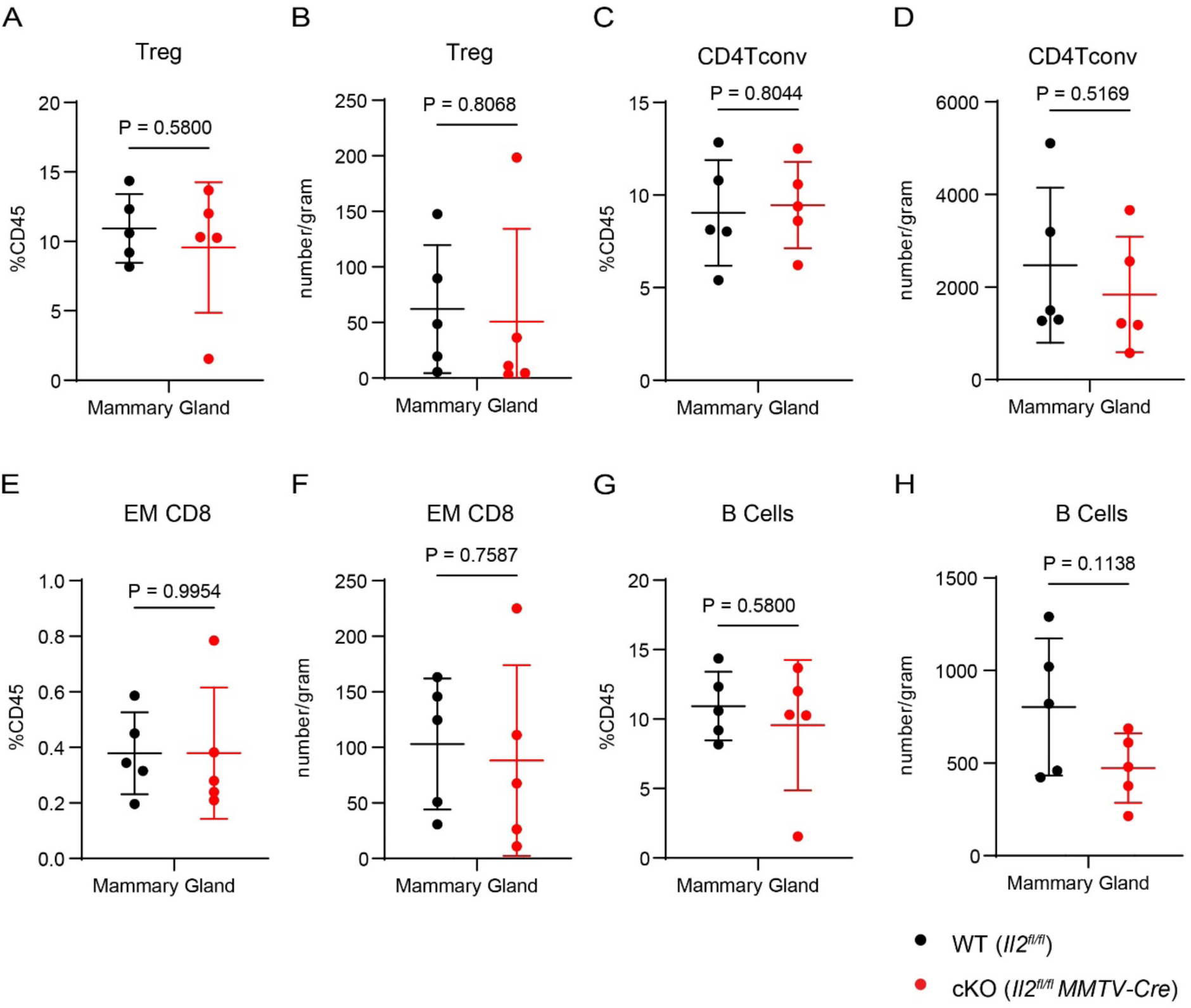
Other immune cell population characterization between WT and MEC-cKO mice. **A.** Percent of Treg cells within the total CD45^+^ cells in the MG. **B.** Treg cell number (per gram) in the MG. **C.** Percent of CD4conv cell of the total CD45^+^ cells in the MG. **D.** CD4conv cell number (per gram) in the MG. **E.** Percent of effector memory CD8 cells of the total CD45^+^ cells in the MG. **F.** Effector memory CD8 cell number (per gram) in the MG. **G.** Percent of the B cells of the total CD45^+^ cells in the MG. **H.** B cell number (per gram) in the MG. Two-sided unpaired *t* test was performed, with P values indicated. NS, not significant. Supplementary Figure for Figure 3.

**Supplementary Figure 6.**
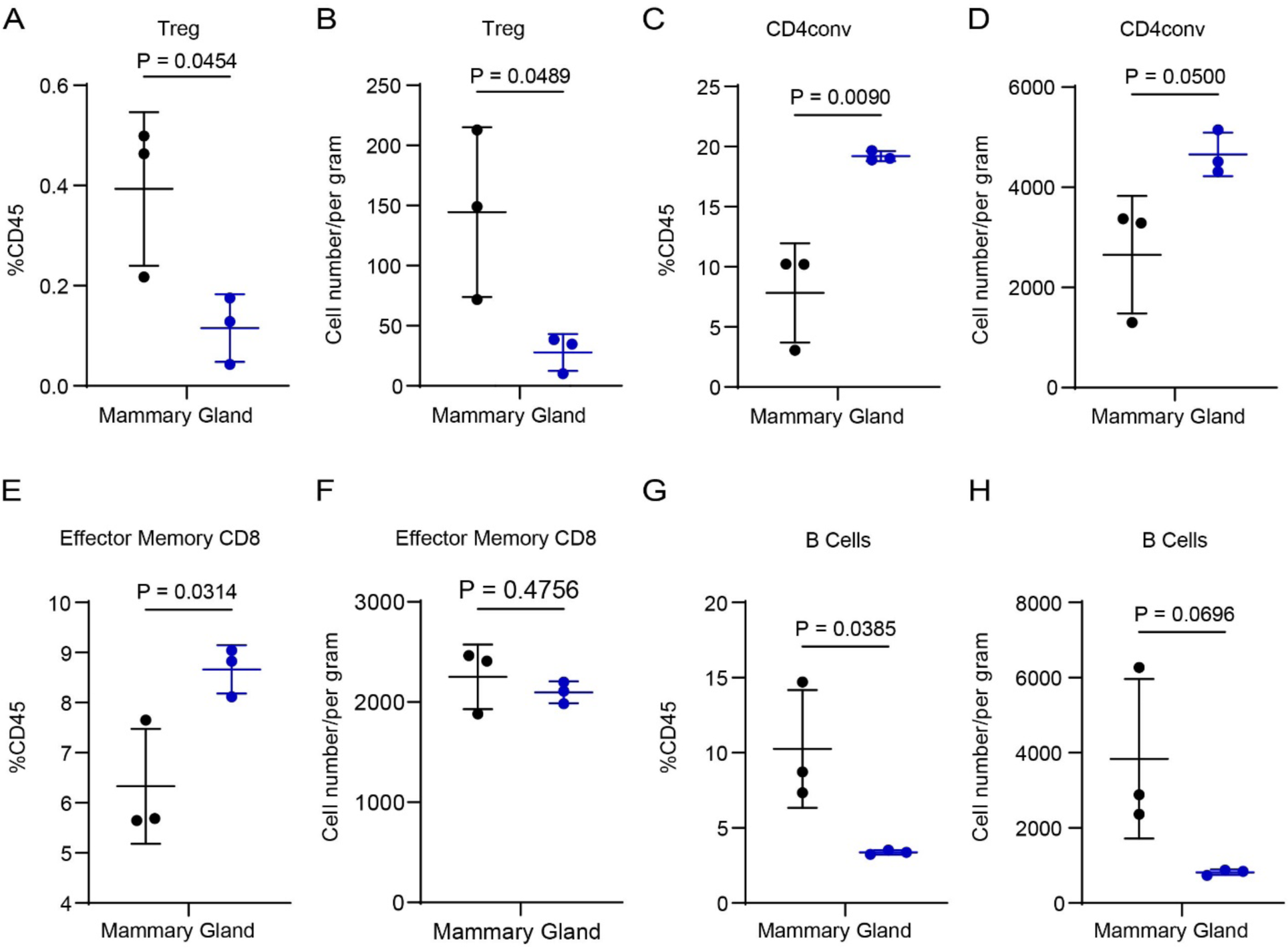
Other immune cell population characterization between WT and T-cKO mice. **A.** Percent of Treg cells within the total CD45^+^ cells in the MG. **B.** Treg cell number (per gram) in the MG. **C.** Percent of CD4conv cell of the total CD45^+^ cells in the MG. **D.** CD4conv cell number (per gram) in the MG. **E.** Percent of effector memory CD8 cells of the total CD45^+^ cells in the MG. **F.** Effector memory CD8 cell number (per gram) in the MG. **G.** Percent of the B cells of the total CD45^+^ cells in the MG. **H.** B cell number (per gram) in the MG. Two-sided unpaired *t* test was performed, with P values indicated. NS, not significant. Supplementary Figure for Figure 4.

**Supplementary Figure 7.**
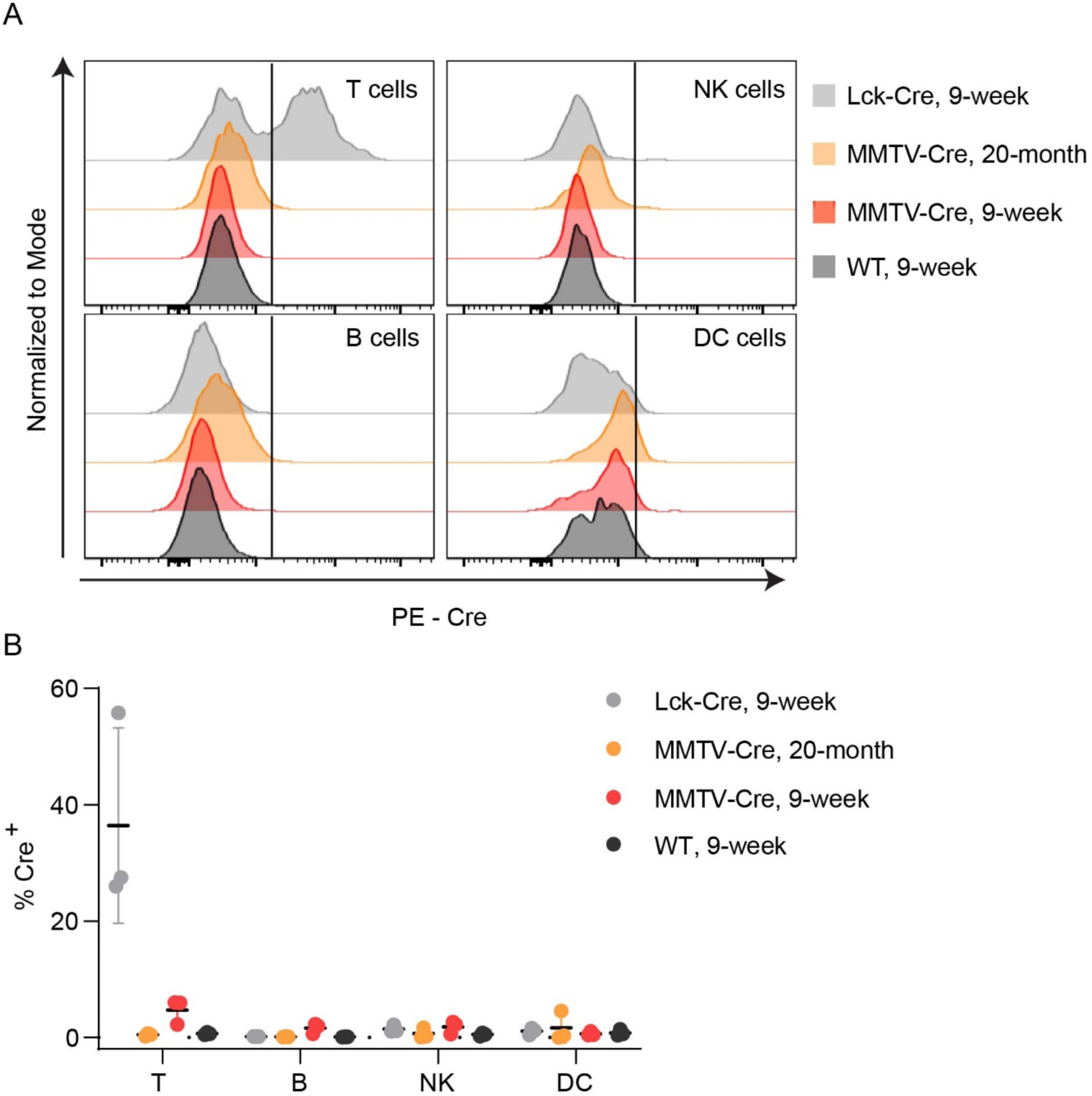
MMTV-Cre expression in different immune cells. **A.** The histogram of Cre (MMTV-Cre and LCK-Cre) in T, B, NK and dendritic cells for flow cytometry analysis between young and old mice. **B.** Quantification of Cre positive percentage in T, B, NK and dendritic cells. Supplementary figure for Figure 3.

**Supplementary Figure 8.**
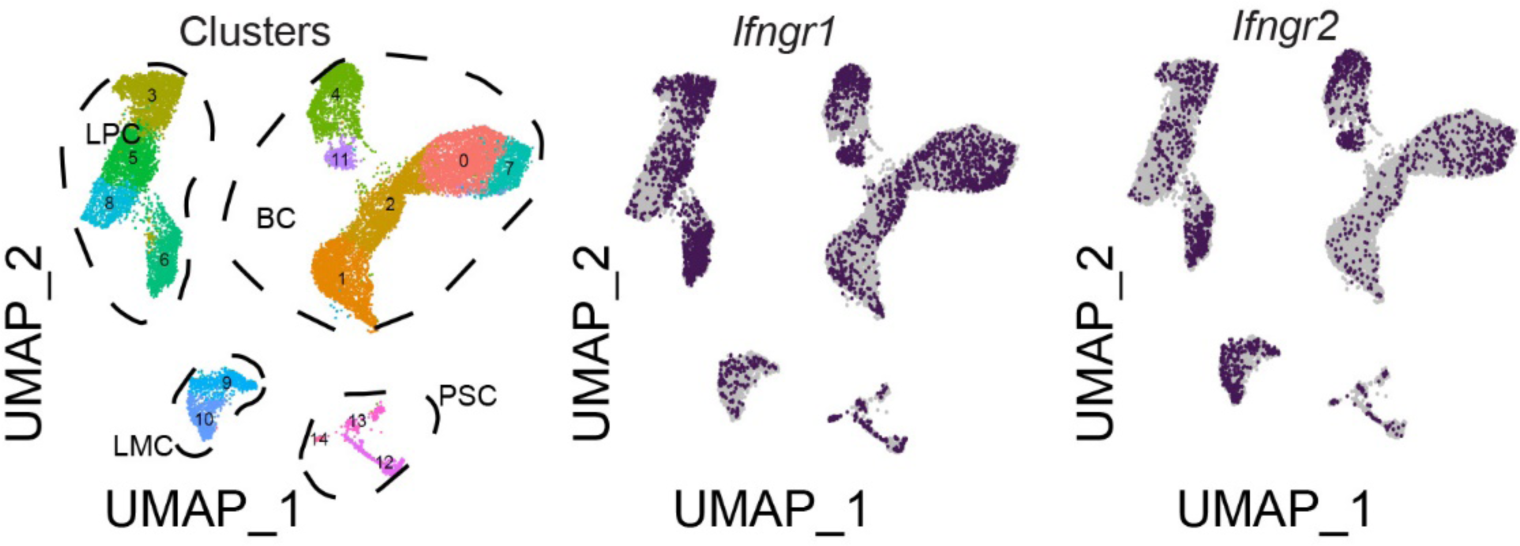
*Ifngr1* and Ifngr2 expression on mouse mammary epithelial cells using single cell RNAseq data from Fig. 1A-1B. Supplementary figure for Figures 5-6.

